# Riboformer: A Deep Learning Framework for Predicting Context-Dependent Translation Dynamics

**DOI:** 10.1101/2023.04.24.538053

**Authors:** Bin Shao, Jiawei Yan, Jing Zhang, Allen R. Buskirk

**Affiliations:** Department of Molecular and Cellular Biology, Harvard University, Cambridge, MA, USA; Department of Chemistry, Stanford University, Stanford, CA, USA; Biological Design Center, Boston University, Boston, MA, USA; Department of Molecular Biology and Genetics, Johns Hopkins University School of Medicine, Baltimore, USA; Klarman Cell Observatory, Broad Institute of Harvard and MIT, Cambridge, MA, USA

## Abstract

Translation elongation is essential for maintaining cellular proteostasis, and alterations in the translational landscape are associated with a range of diseases. Ribosome profiling allows detailed measurement of translation at genome scale. However, it remains unclear how to disentangle biological variations from technical artifacts and identify sequence determinant of translation dysregulation. Here we present Riboformer, a deep learning-based framework for modeling context-dependent changes in translation dynamics. Riboformer leverages the transformer architecture to accurately predict ribosome densities at codon resolution. It corrects experimental artifacts in previously unseen datasets, reveals subtle differences in synonymous codon translation and uncovers a bottleneck in protein synthesis. Further, we show that Riboformer can be combined with *in silico* mutagenesis analysis to identify sequence motifs that contribute to ribosome stalling across various biological contexts, including aging and viral infection. Our tool offers a context-aware and interpretable approach for standardizing ribosome profiling datasets and elucidating the regulatory basis of translation kinetics.

## Introduction

Translation elongation is a critical determinant of protein homeostasis and cellular function^1–3^. Elongation rates across the transcriptome are shaped by a complex interplay between local sequence features, such as mRNA secondary structures, clusters of charged amino acids, and consecutive proline residues, and global factors like cellular resource availability and protein quality control^4–6^. These intricacies impact translation efficiency, co-translational protein folding, and covalent modification^1, 3^. Despite recent advances in understanding translation dynamics, deciphering the regulatory code of ribosome stalling and proteostasis collapse in complex diseases remains challenging^7, 8^.

The advent of ribosome profiling has led to substantial progress in understanding of mRNA translation^5^. Ribosome profiling captures and sequences mRNA fragments protected within ribosomes, allowing the reliable inference of the ribosomal decoding site in each footprint and yielding information about ribosome distribution along mRNA from each gene. Sophisticated models, such as probabilistic models and neural network models, have been used to study ribosome distribution based on mRNA sequences and the biophysical features of the nascent polypeptide^4, 6, 9–12^. However, these studies primarily focus on specific experimental conditions, ignoring the dynamic changes in translation kinetics due to perturbations in cellular states. Whole-cell models provide a precise depiction of the physical process of translation^13, 14^, but fail to account for the bias in ribosome profiling techniques. These limitations result in several consequences: (1) it remains a challenge to distinguish technical artifacts from biological determinants of ribosome elongation, especially when the experimental protocol has a profound effect on the *in vivo* translational landscape^15, 16^; (2) it is hard to identify sequence features that determine the translation kinetics in complex physiological states, which is critical in late on-set misfolding diseases^8^; (3) The predictive power of the model is limited, preventing integrative analysis of large-scale, heterogeneous datasets.

In this paper, we present Riboformer, a deep learning-based framework that models the context-dependent changes in ribosome dynamics at codon resolution. Our approach uses a transformer architecture that detects long-range dependencies in the regulation of elongation^17^ (Fig. 1a). It also utilizes a reference input to prevent the learning of noninformative signals due to the experimental bias. We have benchmarked the performance of Riboformer using a variety of prokaryotic and eukaryotic ribosome profiling datasets. We demonstrate the effectiveness of our network structure in modeling the impact of experimental protocols on the *in vivo* translational landscape, and the trained Riboformer model corrects artifacts in a wide range of unseen datasets. This process reveals subtle differences in synonymous codon translation and uncovers a bottleneck in protein synthesis. Combined with *in silico* mutagenesis analysis, Riboformer identifies peptide motifs that contribute to ribosome stalling across various biological contexts, such as aging and viral infection, highlighting its versatility in diverse research areas (Fig. 1b). Altogether, Riboformer is an end-to-end tool that facilitates the standardization and interpretation of ribosome profiling datasets, and our results demonstrate the potential of context-aware deep learning models, which capture the complex dynamics of biological processes subject to variations in cell physiological states. Riboformer is implemented in Python as a command line tool, publicly available at https://github.com/lingxusb/Riboformer/

**Figure 1.**
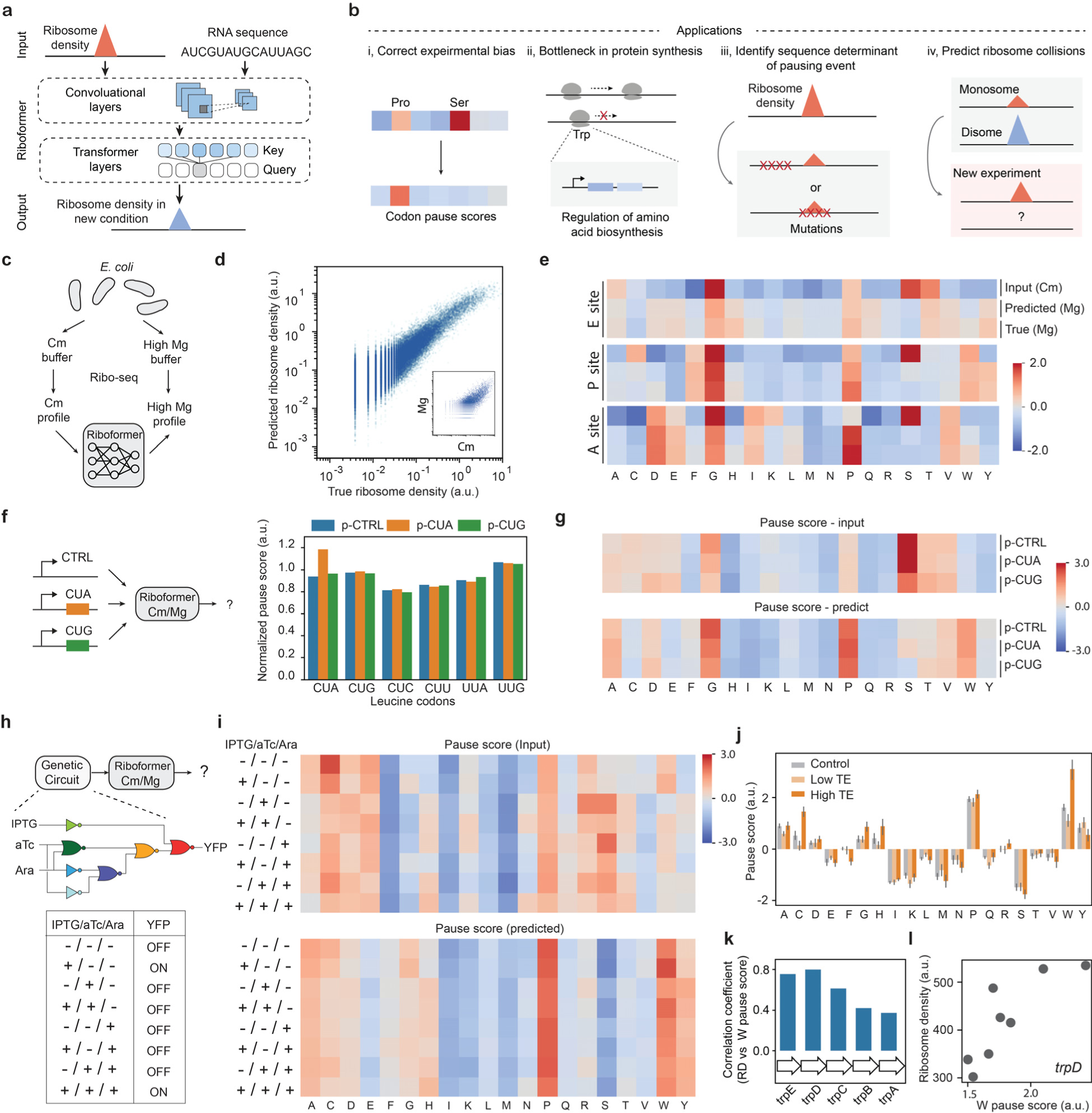
Riboformer captures context-dependency of translation dynamics. **a**, schematic illustration of the neural network model used in the Riboformer pipeline, which is trained to predict ribosome density at single-codon resolution. **b**, applications of the Riboformer pipeline. **c**, *E. coli* cells were treated with different lysis buffers and the resulting profiling data were used to train the neural network model. **d**, correlation between the true and predicted ribosome density for all codon positions with high Mg buffer (Mg) in the test dataset (Pearson correlation coefficient, *r* = 0.91). Inset, ribosome density for all codons with high Mg buffer and Cm buffer (*r* = 0.75). **e**, Heatmap of pause scores for codons for all 20 amino acids. **f**, average ribosome occupancy at leucine codons in endogenous genes after correction of experimental bias. The *E. coli* cells overexpressed a control plasmid (p-CTRL without a mini ORF) or plasmids with a heterologous CUA mini-ORF (p-CUA) or a CUG mini-ORF (p-CUG). **g**, mean codon pause scores for *E. coli* cells overexpressing mini-ORFs. Codon pause scores before (input) and after (predicted) experimental bias correction are shown. **h**, Overview of the procedure to remove experimental artifacts in ribosome profiles from cells with large-scale synthetic circuits. Each gate is associated with a specific transcriptional repressor, which is indicated by the corresponding color^20^. **i**, mean codon pause scores for all eight induction states before and after correction for experiment bias. **j**, pause score of 20 amino acids in genes with different translation efficiency (TE). Error bar represents SEM from eight induction conditions. **k**, correlation of mean Trp pause score with the mean ribosome density (RD) of the operon genes in eight induction states. **l**, *trpD* gene expression and mean Trp codon pause scores in eight induction states (*r* = 0.80).

### Riboformer accurately clarifies ribosome density

Riboformer uses a transformer architecture to capture the context-dependency of translation kinetics (Fig. 1a). The transformer block consists of self-attention layers that gather the impact of distant codons based on their sequence representations^17^, in contrast to convolutional neural network that relies on convolution operators to detect local sequence motifs. As a separate reference input, we also included normalized ribosome density from the control experiment as a baseline for modeling translation dynamics. More specifically, our approach assumes that the relative change in ribosome occupancy is primarily determined by the surrounding sequences. The codon sequence around the position of interest and the normalized ribosome footprint counts in the control experiment were encoded as vectors, which were further connected to two branches of neural networks. The features extracted from the two inputs by a series of transformer blocks were subsequently merged using element-wise multiplication. Finally, a fully connected layer converts the output to the normalized ribosome density in the target condition (methods).

To evaluate the performance of Riboformer, we used bacterial samples that were subjected to perturbations in translation kinetics. Historically, bacterial samples were commonly harvested by rapid filtering and lysed in a buffer containing chloramphenicol (Cm) to arrest elongation^18^. However, recent ribosome profiling and toeprinting studies have found that this protocol alters translation elongation in a sequence-specific manner^16, 19^. To address this issue, a novel protocol was developed that involves flash-freezing the cell culture directly and arresting translation with a lysis buffer containing high magnesium concentrations^16^. This approach eliminates pauses at Ser and Gly codons arising from the filtering protocol and provides a clearer view of the *in vivo* translational landscape. We trained our Riboformer model on this dataset to predict the unperturbed ribosome profile (Mg) based on the perturbed profile (Cm). The input sequence included instances of the codon of interest across all expressing genes (methods) as well as the sequence and ribosome density 20 codons upstream and downstream. The normalized ribosome densities from the rapid filtering experiment were used as the reference input (Fig. 1c). A dataset of 323,688 instances of codons was collected, which was then randomly separated into the training and validation sets.

As shown in Fig. 1d, using samples obtained by filtering with the Cm-lysis buffer, Riboformer accurately predicts the codon-level ribosome density of samples obtained by flash-freezing with the high-Mg buffer. There is a high correlation between the ground truth and the predicted ribosome density (*r* = 0.91, Fig 1d and e). We defined the ratio of ribosome occupancy at each codon to the average ribosome occupancy of the CDS as the codon pause score, and we found that Riboformer recapitulated the average pause score for all the codons (Fig 1f, methods). Notably, ribosome pausing at Gly and Ser codons is largely reduced, and Pro has a high pause score at all the three ribosomal tRNA binding sites (E, P, A) in the corrected profiles^16^. Riboformer also shows superior performance compared to existing models (Supplementary Fig. 1).

### Riboformer corrects experimental bias in unseen data

We further used the trained *E. coli* Riboformer model to correct for the bias in the translational landscapes in other datasets produced with the same experimental artifacts. We applied it to an unseen ribosome profiling dataset obtained by filtering and Cm (Fig. 1g) from *E. coli* cells with low levels of m^1^G37 in tRNAs, a deficiency which affects the decoding of specific codons^20^. Using the trained Riboformer model to predict the unperturbed ribosome occupancy, we were able to correct bias in the pause scores for Gly codons, while maintaining the high pause scores for the affected Pro and Arg codons CCA, CCG, and CGG (Supplementary Fig. 2). Working with a second dataset prepared in a different lab^21^, Riboformer removed the strong pauses at Ser and Gly codons and highlighted increased ribosome occupancy at Pro and Trp codons (Fig. 1h). Moreover, in a sample from this dataset overexpressing a transgene containing the rare Leu codon CUA, we observed a high pause score for the CUA codon in the corrected ribosome profiles, similar to the uncorrected results^21^ (Fig. 1g). Together, these results show that the subtle variation of ribosome pausing in synonymous codons is preserved even as the experimental bias is removed. In addition, the ribosome occupancy from these samples was previously correlated with the level of genome-wide RNA structures determined by dimethyl sulfate (DMS)-seq^22^. Our corrected ribosome occupancy shows a higher correlation with the DMS-seq score (Supplementary Fig. 3) than originally reported^21^, confirming the impact of mRNA secondary structure on translational efficiency^23^. Collectively, these results demonstrate that the Riboformer is a predictive framework that can be used to standardize a wide range of ribosome profiling measurements, reducing experimental noise while remaining true to the underlying biological signal of interest.

### Riboformer allows identification of limiting steps in protein synthesis

In synthetic biology, the proper functioning of engineered systems relies on the coordinated expression of functional genes. However, the expression of heterologous genes imposes an additional burden to the cells, which negatively impacts growth rate and leads to evolutionary instability. Ribosome profiling has been used to quantify the consumption of cellular resources by a 3-input genetic circuit consisting of seven NOT/NOR gates in *E. coli* cells^24^ (Fig. 1h). To gain a better understanding of the translation dynamics in burdened cells, we used the trained Riboformer model to predict the unperturbed ribosome occupancy across the transcriptome in eight circuit states. We found a reduction in the pause scores of Gly, Ser, Thr codons in the ribosomal A site, while Pro and Trp showed the highest pause scores (Fig. 1i). Interestingly, we found that genes with high translational efficiency (TE) tend to have a higher pause score for Trp (Fig. 1j, methods). Trp codons were also enriched in the first 80 codons in the low TE genes (Supplementary Fig. 4). Thus, our results indicate that decoding of Trp codons is a potential rate-limiting step in protein synthesis. To further characterize the role of pausing at Trp codons on gene expression in the strains expressing the engineered circuits, we calculated the correlation of the Trp pause score and the level of expression of the Trp biosynthesis genes for different circuit states, as quantified by the ribosome density (RD) (Fig. 1k). There was a positive correlation between the expression of Trp operon genes and the Trp pause score in the corrected ribosome profiles, especially for *TrpD* and *TrpE*. This observation is in accord with the well-characterized regulation of these genes by transcriptional attenuation after *trpL* which is upstream of *trpE*^25^. Ribosome stalling in the Trp codon-rich *trpL* sequence promotes transcription of the TrpEDCBA operon (Fig. 1l). The clarity in the pausing landscape provided by Riboformer allows us to explain these changes in gene expression driven by overexpression of the circuit components in this example.

### Riboformer identifies sequence determinants of ribosome collisions

Prolonged slowing of translating ribosomes can lead to ribosome collisions, triggering ribosome rescue pathways that promote the degradation of the nascent polypeptide^26–28^. Collided ribosomes form nuclease-resistant disomes during the ribosome profiling protocol because they protect the mRNA at the disome interface. Disome profiling experiments allow the genome-wide detection of collided ribosomes by sequencing the disome-protected mRNA fragments^29–32^. To examine the relationship between ribosome collisions and mRNA sequence features, we used Riboformer to identify the sequence determinants of ribosome collisions in budding yeast (*Saccharomyces cerevisiae*, Fig. 2a). Although the monosome and disome densities show a weak correlation across the genome (Supplementary Fig. 5a, *r* = 0.35), our framework successfully predicts the disome profiles based on monosome occupancy (Supplementary Fig. 5a, *r* = 0.75). For all sites with significant ribosome collisions (n = 11,079, Supplementary Fig. 5b), we used an *in-silico* mutagenesis approach to determine the sequences that contribute to ribosome stalling (Fig. 2a). In brief, a sliding window of the codon sequence was randomly mutated, and the corresponding change in the predicted disome occupancy at the position of interest was defined as the sequence impact score (SIS) for the mutation window (methods).

**Figure 2.**
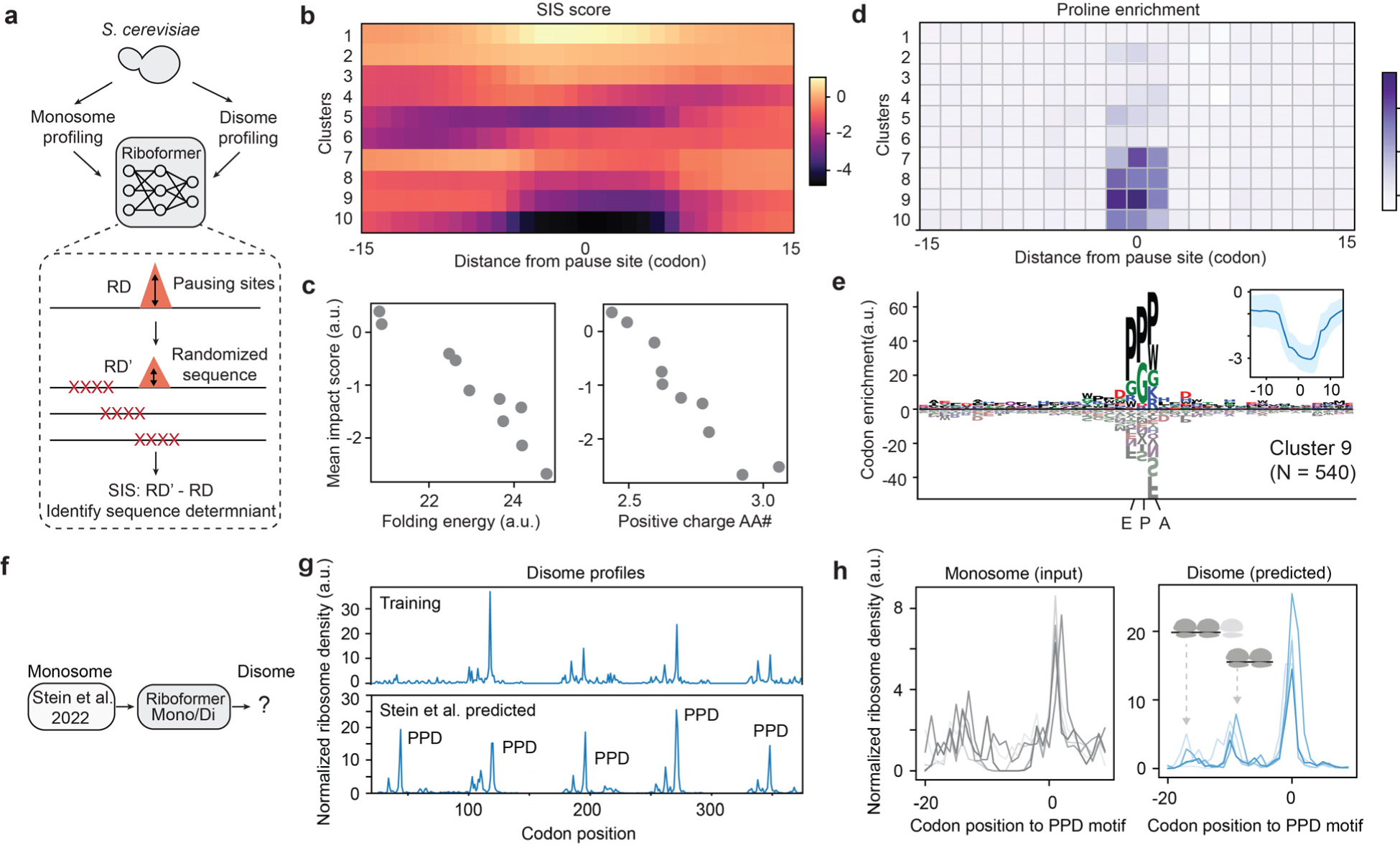
Riboformer identifies sequence determinants of ribosome pausing. **a**, after the Riboformer model is trained, a moving window of codon sequence is randomly mutated and the corresponding change in the predicted ribosome density (dRD) are recorded as the sequence impact score (SIS). **b**, SIS profiles of all the ribosome stalling sites are grouped into 10 clusters. **c**, mean SIS score of each cluster versus the mean folding energy of the input mRNA sequence (left, *r* = −0.96), and the mean number of positive charged amino acids of the input sequence (right, *r* = −0.92). **d**, Pro codon enrichment scores for all the clusters (methods). **e**, codon enrichment profile for cluster 9 (540 ribosome pausing sites). Inset, mean SIS profile for cluster 9. **f**, Trained Riboformer model is used to predict the disome profiles in the Stein et al. dataset^5^. **g**, the monosome and predicted disome profile for gene *UBI4* in the training dataset(top) and Stein et al. dataset (bottom). **h**, the monosome profile before the five PPD ribosome pausing site (left), and the predicted disome profile before the pausing sites (right) are shown for gene *UBI4*.

We performed unsupervised clustering of the SIS profiles for all the significant ribosome collision sites (Fig. 2b, methods) which were grouped into 10 clusters. Interestingly, we found that the mean SIS of each cluster is linearly correlated with the mean mRNA folding energy (*r* = −0.96, Fig 2c). When the mRNA is highly structured, disrupting the mRNA sequence leads to lower levels of ribosome collisions, in agreement with a previous report^33^. Positively charged amino acids have been shown to slow down ribosome elongation speed by interacting with the exit channel^6, 34^. Our approach also identifies a strong negative correlation between the number of positive charged amino acids in the upstream sequence with SIS (*r* = −0.96, Fig. 2c), suggesting that removing these amino acids would reduce ribosome stalling. In contrast, the number of negative charged amino acids show little correlation with SIS (*r* = −0.07, Supplementary Fig. 5c). Notably, a few clusters have their lowest SIS at the ribosome decoding sites (Fig. 2d, clusters 7-10), indicating that the ribosome collisions are mediated by local sequence features. For example, tRNA binding sites in cluster 9 are enriched in Pro (Fig. 2e), which is known to slow down translation elongation^35–37^. In summary, our interpretable framework classifies the ribosome collision events based on their sequence determinants and yields quantitative rules for ribosome stalling.

We further used the trained Riboformer model to identify novel disome sites in yeast from published monosome data^5^ (Fig. 2f). Previous work has demonstrated the regulatory role of ribosome pausing in the processing of ubiquitin peptides. Here we identified five periodic disome peaks in ubiquitin coding gene *UBI4l*, with a novel one at the beginning of the gene, comparing to the training dataset (Fig. 2g). All the peaks were positioned at a proline-rich motif (PPD). When all the disome and monosome profiles are aligned based on the PPD motif, the disome profiles show clear periodic peaks upstream of the pause sites, which is not apparent in the monosome profiles (Fig. 2h).

### Riboformer allows interpretation of exacerbated ribosome stalling in aging

High levels of ribosome collisions can lead to proteostasis collapse in aged organisms^7^. To investigate the mechanism of aging-related ribosome pausing, we applied the Riboformer pipeline on the ribosome profiling data from young and old yeast cells^7^. Using ribosome profiles in young yeast (day 1) as the control, our pipeline successfully predicted ribosome occupancy in aged yeast (day 4, *r* = 0.94, Supplementary Fig. 6a-b). *In silico* mutagenesis analysis of the aging-related pausing sites (n = 6,347 sites, Supplementary Fig. 6c) identified a few clusters with a low SIS at the ribosome decoding site. Further examination of these clusters revealed significant enrichment of Pro codons in the ribosomal E and P site (Supplementary Fig. 6d-f). This observation was not discernible upon analysis of all the ribosome pausing sites^5^. We further extended our analysis to the aging experiments in worms (*Caenorhabditis elegans*, Supplementary Fig. 7). Interestingly, when we examined SIS for the age-dependent pause sites (n = 8,376 sites, Supplementary Fig. 7c), there was an enrichment of Asp codon in the P site for the clusters with similar shapes (Supplementary Fig. 7d-f). In both aged yeast and worm, the SIS score was *positively* correlated with the number of positively charged amino acids (Supplementary Fig. 6h and 7h), unlike what we observed with yeast disomes. Interestingly, the overloaded RQC pathway in aging organisms doesn’t target highly positively charged protein sequences^38^, which may explain the observed correlation.

In our analyses of yeast disomes described above, we observed a negative correlation between mRNA folding energy and SIS, indicating that the ribosomes are more likely to pause in structured regions of mRNA. This correlation holds true for predicted ribosome density from day 4 yeast cells (Supplementary Fig. 6g, *r* = −0.84). Surprisingly, SIS and mRNA folding energy were *positively* correlated in the ribosome collision sites in aged worms (Supplementary Fig. 7g, *r* = 0.97). Our results imply that mRNA secondary structures might play different roles in aging-related ribosome stalling events in these model organisms. Overall, our approach provides a general pipeline for interpretation of context-dependent ribosome pausing and reveals novel insights into how local context affects aging-dependent translation dynamics.

Taken together, our work presents a general predictive framework for standardizing and interpreting ribosome profiling experiments across different organisms and experimental conditions. Our framework models the change in ribosome kinetics caused by the experimental protocol, offering a unique opportunity to correct protocol biases that prohibit the wide adoption of the ribosome profiling technique. We have benchmarked its performance by removing experimental artifacts resulting from rapid filtering and Cm-containing lysis buffer in a range of ribosome profiling datasets. Moreover, by simulating the effect of sequence mutations on ribosome occupancy, the Riboformer model uncovers the impact of amino acid charges and mRNA structure on ribosome collisions and identifies the effect of proline enriched motifs on ribosome stalling in young and aged yeast. This approach provides insight into the regulatory code of translation kinetics, facilitating the discovery of novel therapeutic targets. For example, we applied Riboformer to analyze the ribosome profiles of SARS-CoV-2 following infection of human cells^39^. Our findings reveal that binding motifs of fragile X mental retardation protein (FMRP) contribute to the increased ribosome occupancy in later stages of infection (Supplementary Fig. 8). Notably, FMRP has been demonstrated to bind to polyribosome^40^, and our observation implies the therapeutic potential of fragile X drugs for inhibiting SARS-CoV-2 viral reproduction.

Our Riboformer model provides a means for the integrated analysis of existing ribosome profiling datasets, which are usually generated using different protocols. Comparison of ribosome profiles across multiple species allows the study of ribosome stalling through the lens of evolution, paving the way to investigate the evolutionary forces that determine codon selection and translation elongation efficiency. Further, with the rapid development of single-cell sequencing methods such as single-cell Ribo-seq and TEMPOmap^41, 42^, context-aware models like Riboformer will make it possible to study translation dynamics in a cell state and cell type specific manner. Riboformer can be used as a pure sequence-based model when the reference input is masked, or in combination with other computational methods such as *choros*^43^ to enable more accurate estimation of ribosome distribution. While primarily developed for the ribosome profiling datasets, we envision the Riboformer pipeline could be widely applicable for modeling the experimental bias and biological variations in other types of high-throughput sequencing data.

## Methods

### Ribosome profiling datasets

The ribosome profiling dataset for E coli cells (Cm vs high-Mg lysis buffer) was obtained from NCBI GEO database with accession number GSE119104. The Burkhardt et al., dataset was obtained from NCBI GEO database with accession number GSE77617. The ribosome profiling dataset for genetic circuits was obtained from NCBI GEO database (GSE152664). Genomic data, including gene sequences, as well as transcript and open reading frame (ORF) boundaries, were obtained from NCBI. The *S. cerevisiae* and *C. elegans* aging datasets were downloaded from NCBI GEO (GSE152850). Monosome and disome profiles were obtained from NCBI GEO (GSE139036). The ribosome profiles ofSARS-CoV-2 were obtained from NCBI GEO (GSE149973.). For all the ribosome footprint experiments, we excluded the first and last 10 codons in the downstream analysis to avoid the atypical footprint counts observed at the beginnings and ends of genes. To model ribosome density without being biased by the heterogeneity of translational speed along the 5ʹ ramp and to obtain robust estimates of the steady-state distribution, we excluded all the genes with length < 200 nt. In addition, we filtered out genes with poor ribosome coverage, in accord with previous works^5, 12^. Genes with fewer than 0.5 reads per nucleotide on average in prokaryotes and genes with fewer than 5 reads per nucleotide on average in eukaryotes were excluded from the analysis. For ribosome profiling experiments with replicates, the mean ribosome occupancy at each nucleotide is used for the following analysis. For the codon of interest, we calculated the pause score by taking the mean ribosome density in the 3nt window and dividing it by the mean density across the ORF. The pause scores of codons represent the mean of the scores for all instances of the codon of interest. We further z-score normalized the codon pause scores before visualization.

### Implementation and architecture of Riboformer

We used RNA sequence and the normalized ribosome density in the control experiment as separate input to the Riboformer model. The 120 nt RNA sequence is the input for the first branch. The RNA sequence was converted to the codon sequence and was further transformed into a vector using sequence embedding (hidden dimension: 8). This vector serves as the input of a series of 2D-convolutional blocks, each followed by batch normalization (kernel size: (5,5), filter number: 32, activation function: ReLU). The output of the last 2D-convoluational layer is connected to the transformer layer including a self-attention module and a feed forward module^13^. The transformer block is designed to capture the long-distance dependency of sequences. The self-attention module uses 8 attention heads, and the hidden embedding dimension is 8. The feed forward module uses a 32 node fully connected layer with layer normalization. The last layer in this branch is a 32 node fully connected layer (activation function: ReLU).

The input to the second branch is the ribosome density of the same RNA sequence from the control experiment (120 nt). For each codon, we calculated the sum of reads from all three nt. Then the ribosome density is further log-transformed and processed by a neural network structure that is similar to the first branch. Briefly, it starts with a series of 1D-convolutional blocks (kernel size: (5,1), filter number: 32, activation function: ReLU). Another transformer block follows the last 1D convolutional layer. The output of the second branch has the same dimension as the first branch. Thus, element-wise multiplication was used to combine all the information from the two branches. Finally, a ReLU activation function is used to predict the ribosome density at the position of interest in the new condition.

### Riboformer training and hyperparameter tuning (training and validation dataset construction)

Adam optimizer was used to train the Riboformer model on a A100 GPU (40 GB, Nvidia). A cosine learning decay was used to schedule learning rate with a start learning rate of 0.0005:

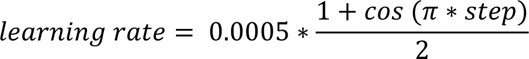

The mean squared error loss function was employed to measure model performance on both the training and validation stages. The explanatory input data and corresponding response variables were divided into training (70%), validation (15%) and test (15%) sets. Early stopping was introduced to prevent overfitting, and the training process terminated when the validation loss did not decrease for 10 epochs. For building and training models, Keras v2.2 and Tensorflow v1.10 software packages were used.

### Supervised shallow methods

Ridge regression model was trained using the sklearn software package (0.24.2). The dataset was randomly divided into the training (70%) and test (30%) sets. One hot encoding was used to convert codon identities to input vectors. Alpha was sampled from 0.001 to 50 and the alpha value with the best performance on the test dataset was selected.

### Codon positional enrichment

We calculated the translation efficiency (TE) for a target gene as the ratio between mean of the ribosome density (RD) and the mRNA expression. The ribosome density (RD) of each gene was calculated by averaging all ribosome occupancies over the length of the gene^20^. The mRNA expression in FPKM (fragments per kilobase of transcript per million mapped reads) of each gene that was calculated by averaging the height of RNA-seq profile over the length of the gene.

We analyzed the first 100 codons of genes with TE in the highest/lowest 10 percentiles among all the genes. In each 10-codon window, we calculated the number of a specific codon. It is then compared with the codon number from a randomly sampled gene group with the same number of genes. We calculated the p values from a student t-test (function *ttest_ind* from the scipy package) as the positional enrichment for the specific codon.

### Identification of conditional-dependent pause sites

To identify conditional-dependent ribosome pausing sites, we used a strategy that is similar to Stein et al.^5^, which utilized a two-tailed Fisher’s exact tests to identify codon positions with statistically significant changes in ribosome pausing. At each codon position, a 2X2 contingency tables were created to perform a two-tailed Fisher’s exact test to compare the ratio of the reads in the control sample and the sample of interest. This compares the observed ratio of ribosome reads at a specific position from the two samples to the expected ratio based on total number of reads from the two samples. It allows the calculation of the odds ratio as well as the p value. The first 10 and last 10 codons of the transcript were excluded in the analysis. The conditional pausing sites were identified as follows: p value < 0.001 and odds ratio > 1.

### *In silico* mutagenesis analysis

For each conditional dependent pausing site, we denote the Riboformer predicted ribosome density as RD. In the 40-codon input sequence, we selected a 10-codon window and sampled 100 random sequences {*x_j_*} to replace the original sequence. The mean predicted ribosome density from the random sequences was calculated as 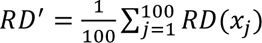 and RD – RD’ is the sequence impact score (SIS) for the 10-codon window. We moved the window along the RNA sequence at one codon step so that every codon is randomly mutated 1000 times. Enrichment of known sequence motifs of RNA-binding proteins were identified using SEA^44^.

### Clustering analysis of sequence impact scores

We used the K-means clustering method from the Python scikit-learn package to cluster the impact score profiles. Elbow method was used to determine the cluster number and the random seed was set to 0.

For each 120 nt RNA sequence, we calculated its folding energy using the RNAfold software (https://www.tbi.univie.ac.at/) with default parameters. The energy was then averaged for each cluster.

To calculate the codon enrichment for each cluster, the codon occurrences at each position (−20 to 20) for each cluster were compared with randomly sampled codon sequences. A student t test was used to calculate the p value of the enrichment or depletion of the specific codons. The sequence log was generated based on the log-transformed p values.

## Data availability

We provide all datasets generated or analyzed during this study. The ribosome profiles were downloaded from Gene Expression Omnibus with the accession numbers GSE119104 (Mohammad dataset^12^), GSE77617 (Burkhardt dataset^16^), GSE98664 (synthetic circuit dataset^20^), GSE152850 (aging dataset^21^), GSE139036 (disome dataset^5^), GSE149973 (SARS-CoV-2 dataset^39^). More information for these datasets could be found in Methods.

## Code availability

Code used for training models and performing analyses are available from GitHub (https://github.com/lingxusb/Riboformer)

## Acknowledgement

We are grateful to Y. Yan, R. Majovski, H. Kang and N. Guydosh for discussions. A.R.B was supported by NIH grant GM136960.

## Author contributions

B.S. conceived the research project. B.S. preprocessed raw data, designed the neural network model. B.S. and J.Y. implemented the model and carried out model training and validation tasks. B.S. performed the computational and statistical analyses. B.S., J.Y., J.Z., and A.R.B. wrote the manuscript. All the authors discussed the results and commented on the manuscript.

## Supplementary figures

**Supplementary Figure 1:**
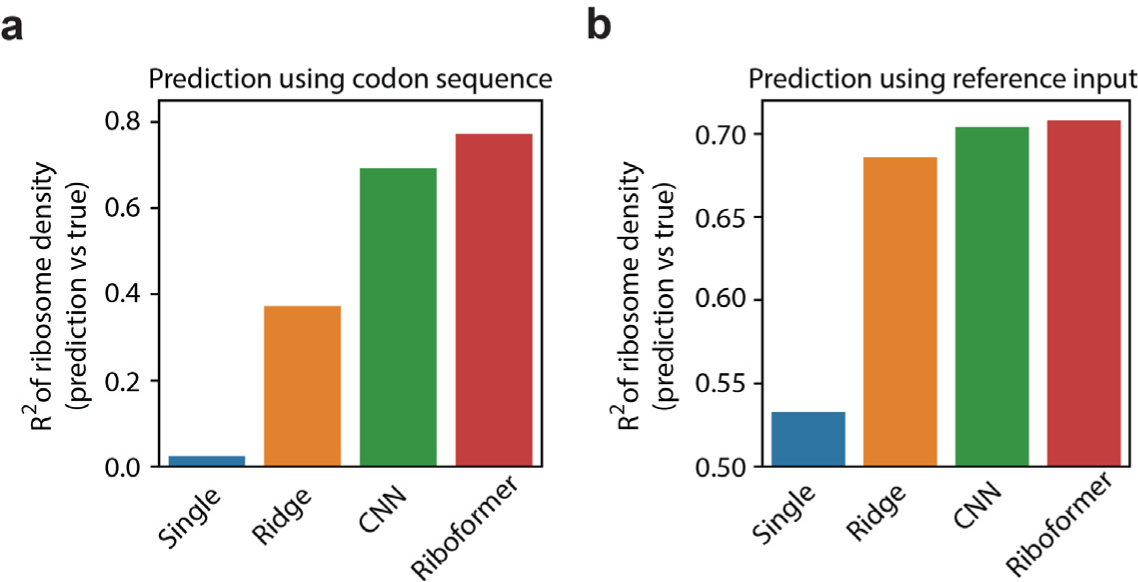
Performance of Riboformer. Bar plot of coefficient of determination (R2) between the true and predicted ribosome density for all codons in the test dataset using both codon sequence (**a**) and the ribosome density in the control experiment (**b**). Ribosome density is predicted using the codon identity (single), ridge regression (ridge), convolutional neural networks (CNN) and Riboformer model. CNN model is the Riboformer model without the transformer layers.

**Supplementary Figure 2:**
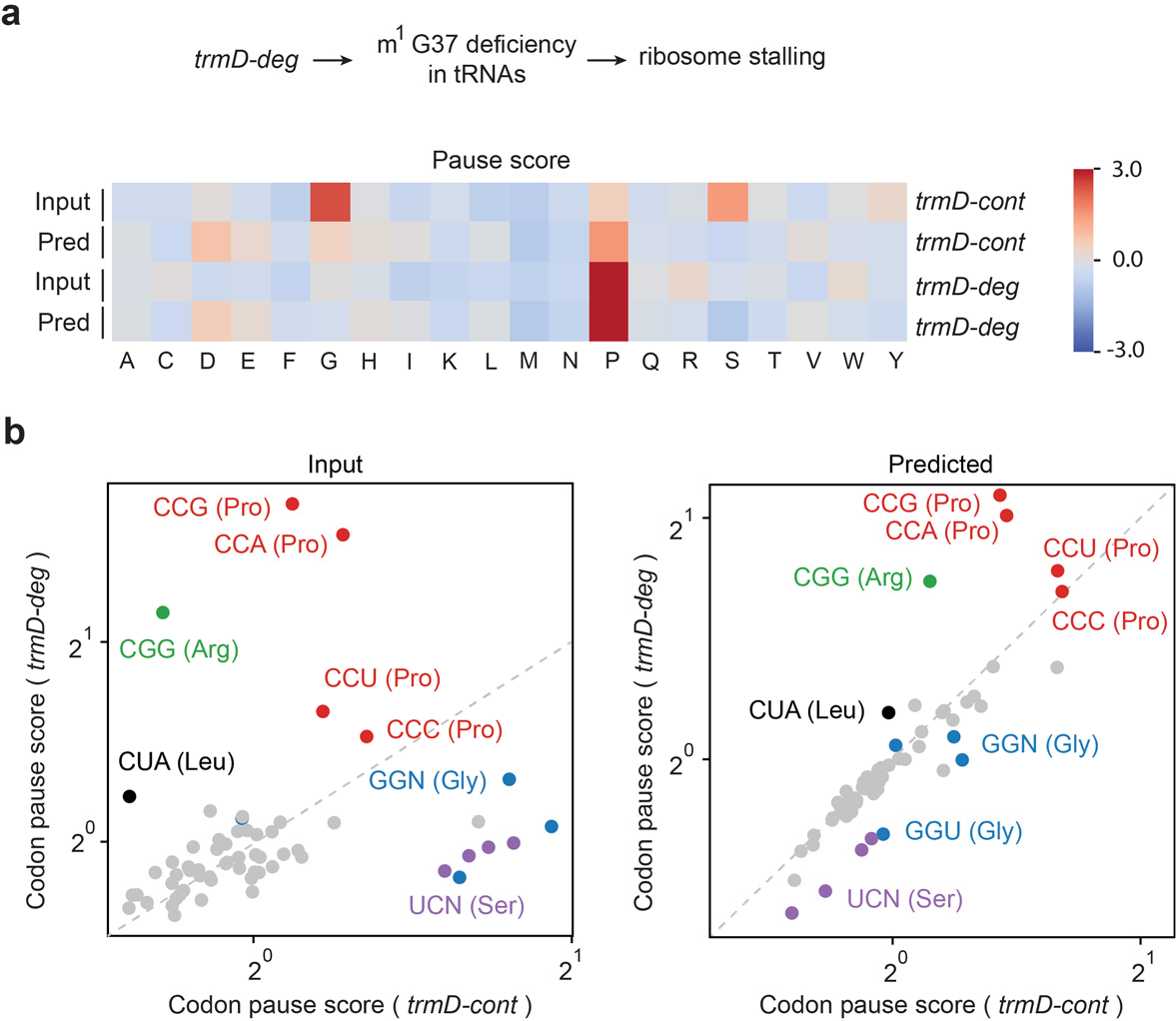
Ribosome pausing in *E. coli* cells with m^1^G37 deficiency. **a**, mean codon pause scores are shown for the WT (*trmD-cont*) and trmD depletion cells (*trmD-deg*). **b**, pause scores for codons positioned in the ribosomal A site before (left) and after (right) correction for the experimental bias. 61 sense codons are shown individually. Riboformer was used to remove the experimental bias in the ribosome profiles.

**Supplementary Figure 3:**
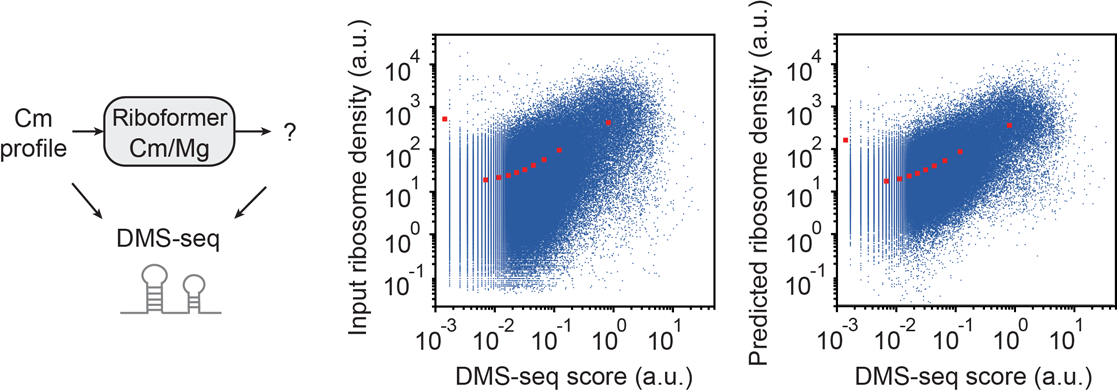
Codon level comparison of ribosome density and DMS-seq. **a,** Trained Riboformer model was used to correct the ribosome profiles in *E. coli***. b**, codon-level comparison of ribosome density and DMS-seq score before (left, *r* = 0.26) and after (right, *r* = 0.32) correction. Each dot is one codon. Binned average of ribosome densities is shown in red squares.

**Supplementary Figure 4:**
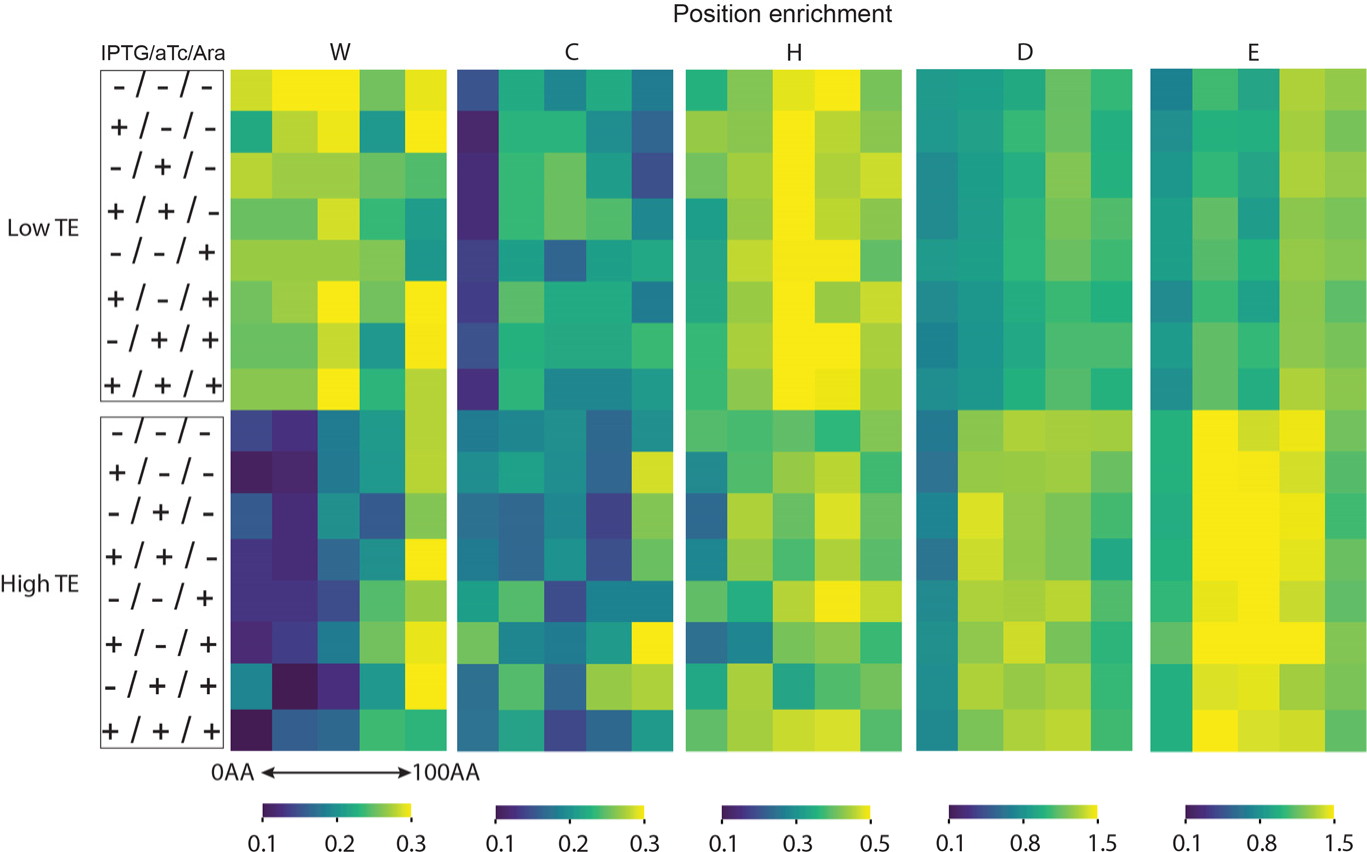
Position enrichment of codons for the low and high TE genes in *E. coli* cells. Low TE and high TE genes are the genes with the top 10% and lowest 10% translational efficiency. Genes that are less than 100 amino acids long were filtered out.

**Supplementary Figure 5:**
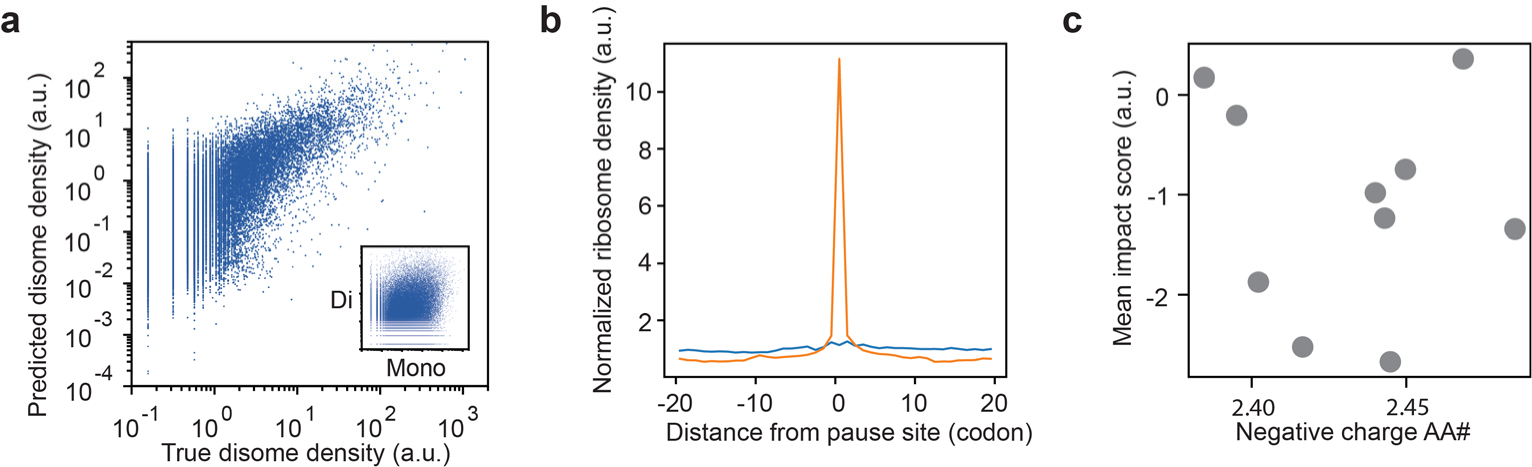
Sequence determinant of disome peaks in yeast. **a,** monosome and disome profiling dataset is used to train the Riboformer model. The predicted disome density and the true disome density in the test dataset is shown (*r* = 0.75). Each dot is a single codon of interest. Inset, comparison of disome density and monosome density (*r* = 0.35). **b,** average ribosome occupancy of the disome profiles around disome formation sites (orange) and the average ribosome occupancy in the monosome profiles (blue) are shown, n = 11,079 sites. **c**, Comparison between the number of negative charged amino acids and the SIS in yeast (*r* = −0.07). Each dot represents mean number of the negative charged amino acids in the input sequence and the mean SIS for one cluster (Fig. 2c).

**Supplementary Figure 6:**
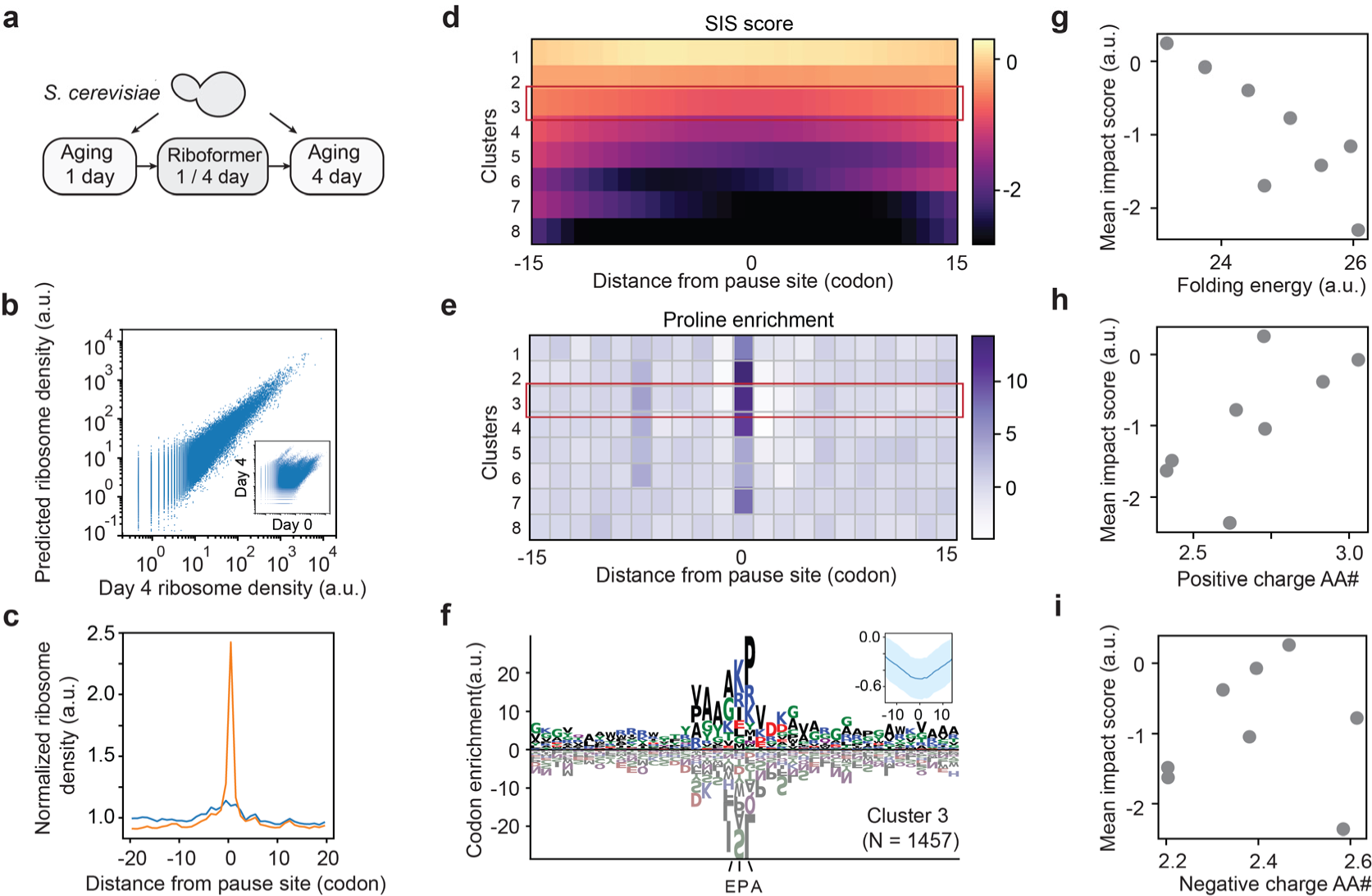
Sequence determinant of ribosome pausing in aged yeast. **a**, ribosome profiles from young and aged yeast were used to train the Riboformer model. **b**, the predicted ribosome density and true ribosome density in the test dataset is shown. Each dot is a single codon of interest, *r* = 0.94. Inset, comparison of ribosome density in young and old yeast cells, *r* = 0.44. **c**, the average ribosome occupancy at age-dependent pause sites. n = 6,347 sites. **d**, SIS profiles of all the ribosome stalling sites are grouped into 8 clusters. Cluster 3 is highlighted, which has the lowest SIS score in the decoding site. **e**, proline codon enrichment scores for all the clusters (methods). **f**, codon enrichment profile for cluster 3 (1,457 ribosome pausing sites). Inset, mean SIS profile for cluster 3. g-i, comparison of the folding energy of the input sequence, *r* = −0.84 (**g**), mean number of positive charged amino acid, *r* = 0.68 (**h**), mean number of the negatively charged amino acid, *r* = 0.05 (**i**) of the input sequence with the mean SIS for 8 clusters. Each dot represents mean values from one cluster.

**Supplementary Figure 7:**
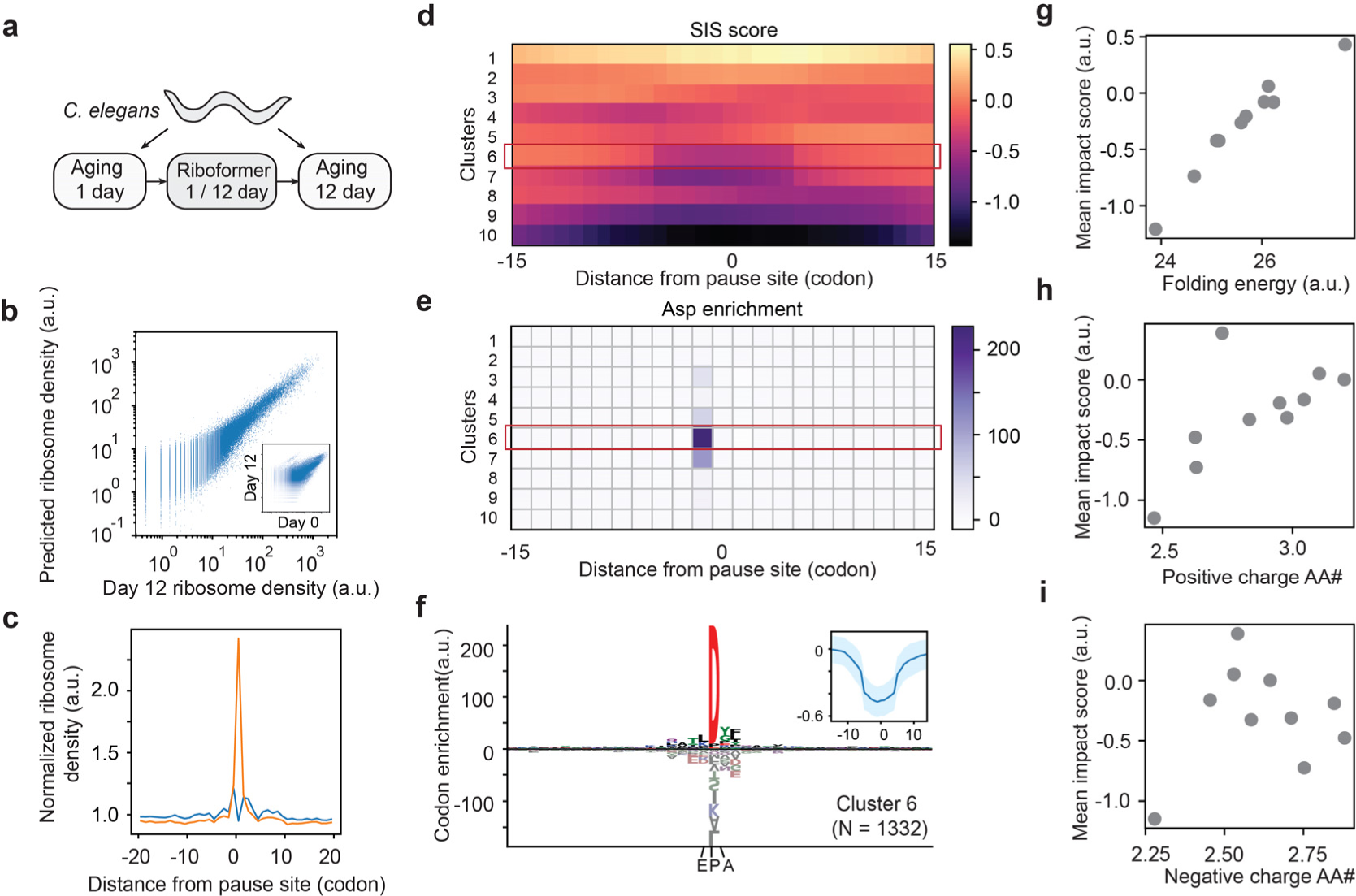
Sequence determinant of ribosome pausing in aged worms. **a**, ribosome profiles from young and aged worm cells were used to train the Riboformer model. **b**, the predicted ribosome density and true ribosome density in the test dataset is shown. Each dot is a single codon, *r* = 0.91. Inset, comparison of ribosome density in young and old worms, *r* = 0.81. **c**, the average ribosome occupancy at age-dependent pause sites. n = 8,376 sites. **d**, SIS profiles of all the ribosome stalling sites are grouped into 10 clusters. Cluster 6 is highlighted, which has the lowest SIS score in the decoding site. **e**, Asp codon enrichment scores for all the clusters (methods). **f**, codon enrichment profile for cluster 6 (1,332 pausing sites). Inset, mean SIS profile for cluster 6. g-i, comparison of the folding energy of the input sequence, *r* = 0.97 (**g**), mean number of positive charged amino acid, *r* = 0.66 (**h**), mean number of the negatively charged amino acid, *r* = 0.15 (**i**) of the input sequences with the mean SIS for 10 clusters. Each dot represents mean values from one cluster.

**Supplementary Figure 8:**
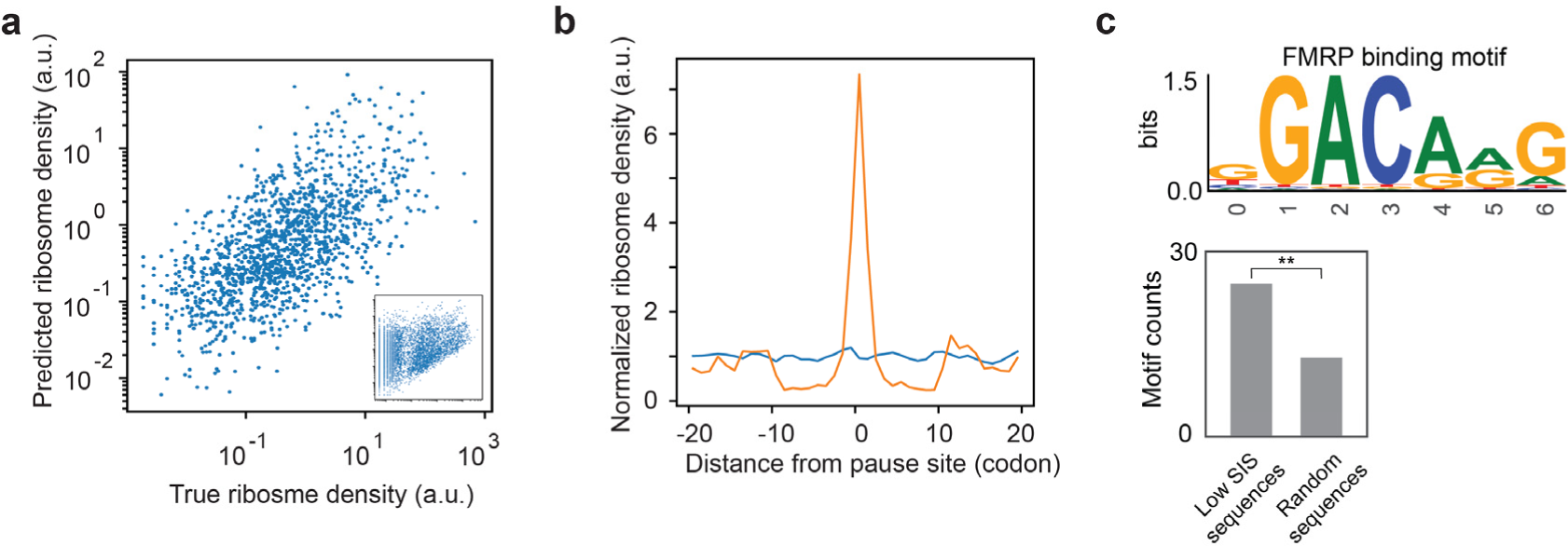
Analysis of the ribosome profiles of SARS-CoV-2 canonical open reading frames (ORFs). **a,** ribosome densities of the SARS-CoV-2 canonical ORFs at 5 and 24 hours post-infection (hpi) in human Vero E6 cells were used to train the Riboformer model. The predicted ribosome density and the true ribosome density in the test dataset at 24 hpi is shown (*r* = 0.63). Each dot is a single codon of interest. Inset, comparison of ribosome densities at 5 hpi and 24 hpi (*r* = 0.43). **b,** the average ribosome occupancy of the 24 hpi profiles around the corresponding ribosome stalling sites (orange) and the average ribosome occupancy in the 5 hpi profiles (blue) are shown, n = 170 sites. **c**, upper panel: binding motif of FMRP; lower panel: Comparison of the number of FMRP binding motifs in the sequences with low SIS (below −3, 10 amino acids in length, n = 65) versus the number of FMRP binding motifs in an equal quantity of random sequences with identical length. **p-value = 0.017.

